## Supplementary figures and images for "Empirical examination of the replicability of associations between brain structure and psychological variables"

### Figure S1

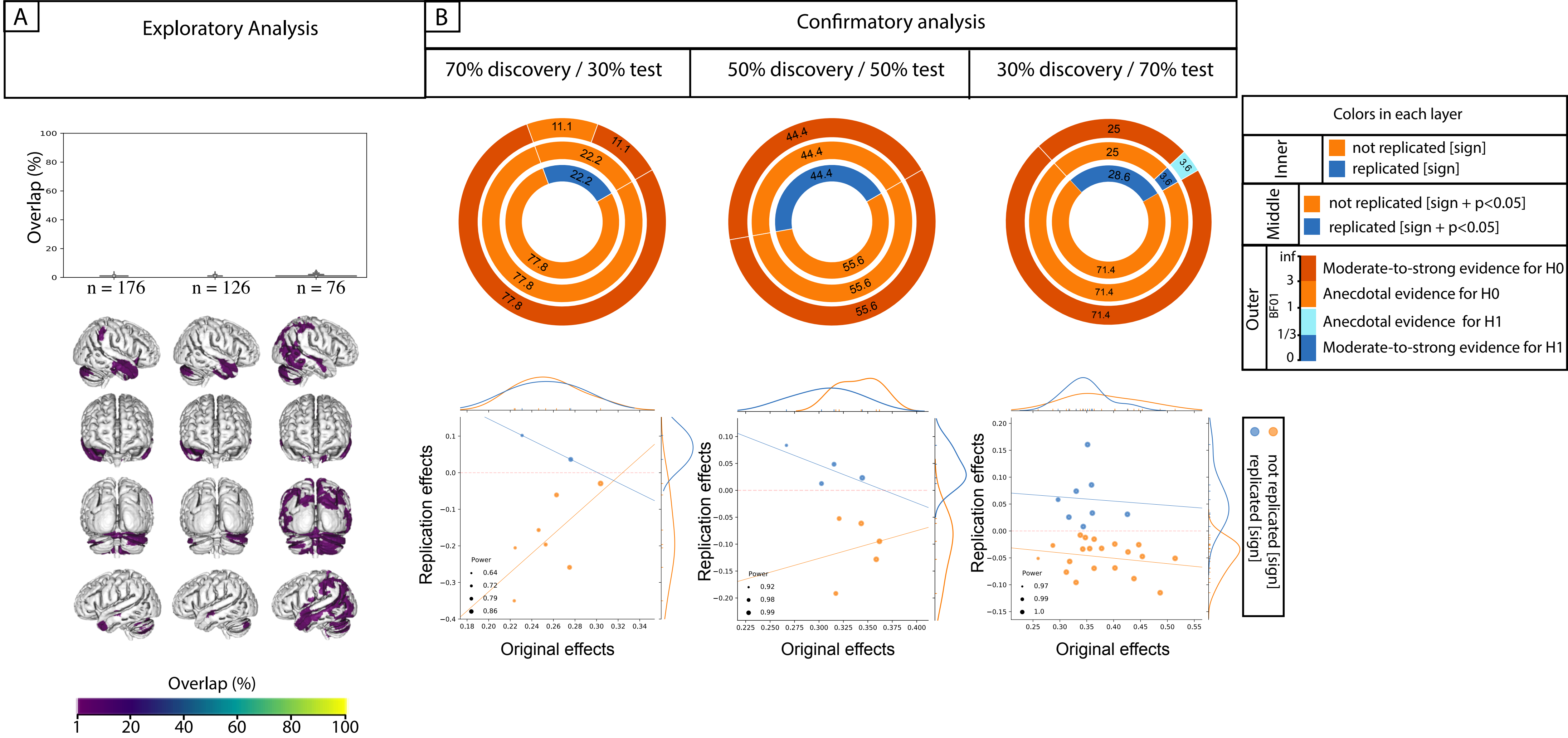

### Figure S2

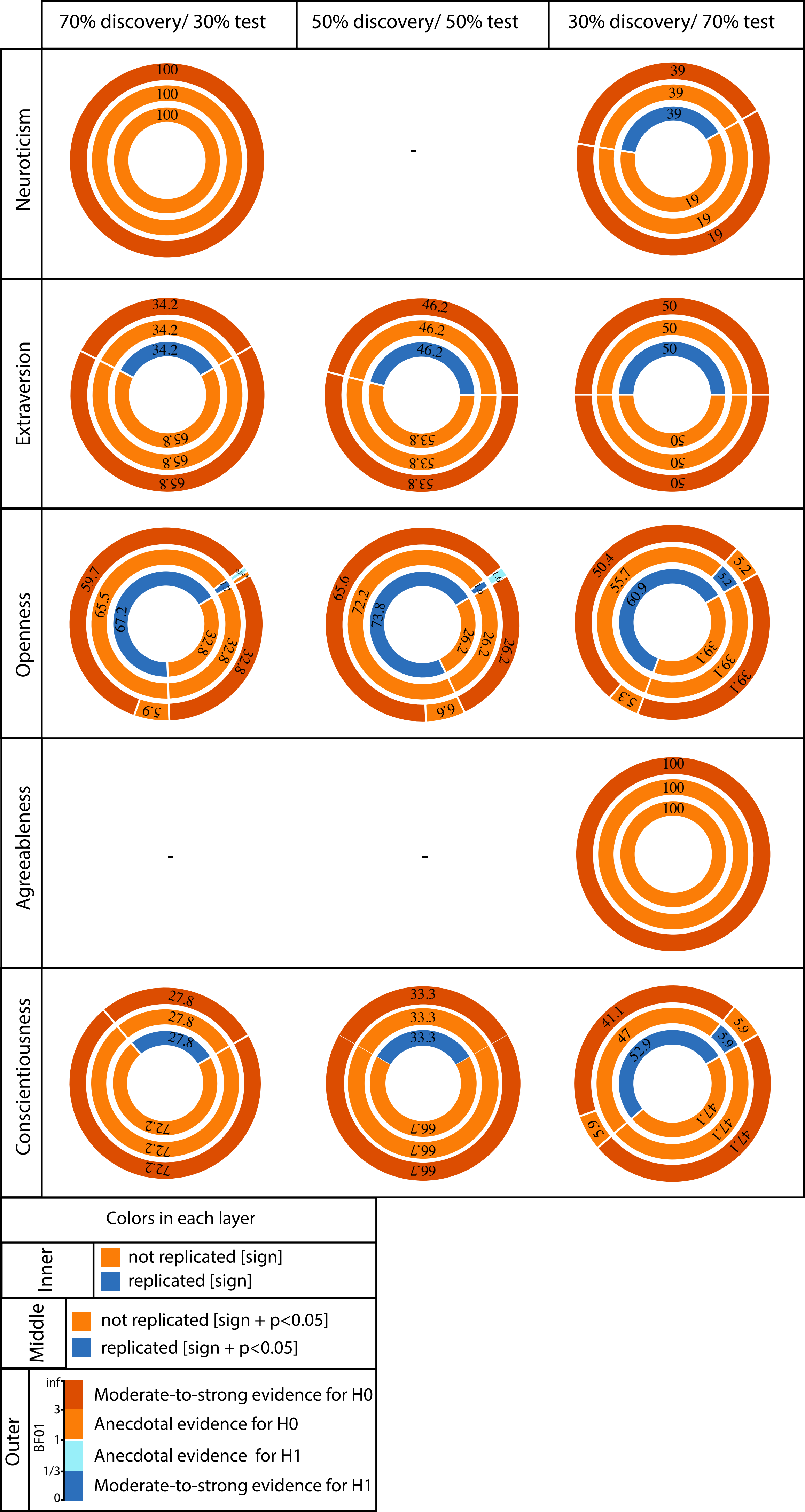
