## Supplementary Tables for "Empirical examination of the replicability of associations between brain structure and psychological variables"

**Table S1.** Distribution of the raw phenotypical and psychological scores in the whole sample.

| **Healthy sample** | **Participants** (n = 466 ; 153 male) |
| --- | --- |
| Age (years)  n-total = 466 | 48.34 $\pm$19.2 (18 , 85) |
| BMI (kg/m^2^)  n-total = 466 | 27.6 $\pm$5.8 (16 , 56) |
| Education (years)  n_total = 466 | 15.5 $\pm$2.25 (10 , 24) |
| Card Sorting (Free sorting)  n-total = 259 | 37.8 $\pm$10.8 (8 , 63) |
| Card Sorting (Sort recognition)  n-total = 258 | 35.5 $\pm$12.37 (8 , 63) |
| Card Sorting (sort recognition versus free sort)  n-total = 257 | -0.76 $\pm$2.1 (-7 , 4) |
| Total IQ (sum of t-scores)  n-total = 466 | 204.1 $\pm$28.7 (95 , 289) |
| Perceptual IQ (sum of t-scores)  n-total = 466 | 100.1 $\pm$17.2 (54 , 170) |
| Vocabulary IQ (sum of t-scores)  n-total = 466 | 104.3 $\pm$15.68 (21 , 149) |
| ANT (Alert) (msec)  n-total = 455 | 34.4 $\pm$31 (-88,145) |
| ANT (Orient) (msec)  n-total = 449 | 19.4 $\pm$20.2 (-44 , 90) |
| TMT (Visual scanning) (sec)  n-total = 457 | 20.5 $\pm$5 (10 , 37) |
| TMT (number sequencing) (sec)  n-total = 452 | 30.5 $\pm$10.6 (11 , 67) |
| TMT (Letter sequencing) (sec)  n-total = 452 | 30.1 $\pm$10.8 (11 , 80) |
| TMT (number-letter switching) (sec)  n-total = 447 | 77.5 $\pm$29.7 (25 , 188) |
| TMT (motor speed) (sec)  n-total = 456 | 26.3 $\pm$8.9 (2 , 58) |
| CWI (color naming) (sec)  n-total = 455 | 29 $\pm$5.6 (17 , 46) |
| CWI (word reading) (sec)  n-total = 450 | 21.5 $\pm$4 (12 ,34) |
| CWI (interference) (sec)  n-total = 449 | 54.8 $\pm$12.7 (26 , 99) |
| CWI (Inhibition/switching) (sec)  n-total = 454 | 62 $\pm$16.9 (32 , 140) |
| Verbal Fluency (letter)  n-total = 453 | 40 $\pm$12 (9 ,77) |
| Proverbs (Free inquiry)  n-total = 356 | 10.8 $\pm$2.5 (3 ,16) |
| Word-context (number of consecutively correct)  n-total = 262 | 25 $\pm$6.1 (7 , 38) |
| 20-questions (initial abstractions)  n-total = 360 | 30.4 $\pm$13.3 (2 , 60) |
| 20-questions (total asked questions)  n-total = 354 | 27.8 $\pm$7 (17 , 55) |
| 20-questions (total weighted achievement score)  n-total = 356 | 15 $\pm$2.7 (6 , 20) |
| RAVLT (T1-immediade recall)  n-total = 254 | 6 $\pm$1.6 (1 , 10) |
| RAVLT (T5-immediade recall)  n-total = 255 | 11.7 $\pm$2.3 (5 , 15) |
| RAVLT (total immediate recall)  n-total = 254 | 48.1 $\pm$9.2 (18 , 67) |
| RAVLT (delayed recall)  n-total = 255 | 9 $\pm$3.5 (0 , 15) |
| RAVLT (delayed correct recognition)  n-total = 251 | 12.7 $\pm$1.9 (6 , 15) |
| RAVLT (delayed false recognition)  n-total = 252 | 1.5 $\pm$1.8 (0 , 8) |
| Anxiety (State)  n-total = 466 | 30.6 $\pm$9.3 (20 , 71) |
| Anxiety (Trait)  n-total = 466 | 33.7 $\pm$9.4 (20 , 67) |
| NEO (N total)  n-total = 462 | 17.2 $\pm$7.4 (0 , 42) |
| NEO (E total)  n-total = 461 | 30.3 $\pm$5.8 (12 , 48) |
| NEO (O total)  n-total = 462 | 32 $\pm$5.9 (16 , 48) |
| NEO (A total)  n-total = 463 | 35 $\pm$5.7 (18 , 48) |
| NEO (C total)  n-total = 463 | 34.6 $\pm$7 (13 , 48) |
| **clinical sample** | **Participants** (n= 371, 200 male; 39 Sites) |
| Age (years) | 71.6 $\pm$7.4 (55 , 91) |
| Education (years) | 16.2 $\pm$2.5 (9 , 20) |
| Diagnosis [n] [SMC / EMCI / LMCI / AD] | 47 / 177 / 85 / 62 |
| MMSE | 28 (19 , 30)* |
| RAVLT (total immediate recall) | 36.6 $\pm$12. 75 (1 , 68) |

**Data are mean ± standard deviation (minimum-maximum), unless indicated otherwise.**

*** median (minimum-maximum).**

Abbreviations: BMI : body mass index; IQ : intelligence quotient; ANT : attention network task; CWI: color-word interference task; RAVLT : Rey auditory verbal learning task; NEO five factor inventory (N: neuroticism, E: extraversion, O: openness, A: agreeableness, C: conscientiousness); SMC: significant memory complaint; EMCI: early mild cognitive impairment; LMCI: late mild cognitive impairment; AD : Alzheimer’s disease.

**Table S2. Summary of the exploratory findings.** For each discovery sample size, the number of clusters in which gray matter volume is positively or negatively associated with the tested psychological score is reported. Number of splits (out of 100) in which the clusters were detected are noted in parentheses.

|  | n_discovery = 70% n_total | | n_discovery = 50% n_total | | n_discovery = 30% n_total | |
| --- | --- | --- | --- | --- | --- | --- |
|  | # positive clusters (%split) | # negative clusters (%split) | # positive clusters (%split) | # negative clusters (%split) | # positive clusters (%split) | # negative clusters (%split) |
| Card Sorting (Free sorting)  n-total = 259 | 67 (17%) | 0 | 55 (13%) | 0 | 59 (15%) | 1 (1%) |
| Card Sorting (Sort recognition)  n-total = 258 | 34 (12%) | 0 | 62 (17%) | 0 | 59 (15%) | 0 |
| Card Sorting (sort recognition versus free sort)  n-total = 257 | 1 (1%) | 0 | 1 (1%) | 0 | 1 (1%) | 3 (2%) |
| Total IQ (sum of t-scores)  n-total = 466 | 137 (27%) | 0 | 128 (27%) | 0 | 56 (16%) | 0 |
| Vocabulary IQ (sum of t-scores)  n-total = 466 | 0 | 1 (1%) | 0 | 2 (2%) | 19 (4%) | 1 (1%) |
| ANT (Alert)  n-total = 455 | 41 (14%) | 0 | 47 (11%) | 0 | 92 (15%) | 0 |
| ANT (Orient)  n-total = 449 | 0 | 0 | 1 (1%) | 0 | 1 (1%) | 13 (3%) |
| TMT (Visual scanning)  n-total = 457 | 0 | 2 (2%) | 0 | 17 (4%) | 0 | 5 (2%) |
| TMT (number sequencing)  n-total = 452 | 1 (1%) | 0 | 4 (2%) | 15 (5%) | 5 (3%) | 38 (6%) |
| TMT (Letter sequencing)  n-total = 452 | 0 | 40 (15%) | 0 | 51 (18%) | 2 (1%) | 42 (9%) |
| TMT (number-letter switching)  n-total = 447 | 0 | 15 (11%) | 0 | 31 (16%) | 0 | 24 (9%) |
| TMT (motor speed)  n-total = 456 | 0 | 21 (9%) | 1 (1%) | 26 (8%) | 0 | 38 (10%) |
| CWI (color naming)  n-total = 455 | 0 | 5 (4%) | 0 | 45 (13%) | 0 | 43 (17%) |
| CWI (word reading)  n-total = 450 | 5 (1%) | 3 (3%) | 3 (3%) | 12 (5%) | 6 (1%) | 5 (3%) |
| CWI (Inhibition/switching)  n-total = 454 | 0 | 85 (25%) | 0 | 113 (21%) | 0 | 72 (19%) |
| Verbal Fluency (letter)  n-total = 453 | 2 (2%) | 0 | 8 (5%) | 0 | 5 (2%) | 1 (1%) |
| Proverbs (Free inquiry)  n-total = 356 | 33 (11%) | 0 | 38 (9%) | 1 (1%) | 66 (18%) | 7 (2%) |
| 20-questions (initial abstractions)  n-total = 360 | 95 (38%) | 0 | 100 (29%) | 0 | 48 (17%) | 0 |
| 20-questions (total asked questions)  n-total = 354 | 0 | 11 (7%) | 0 | 16 (6%) | 2 (2%) | 50 (17%) |
| 20-questions (total weighted achievement score)  n-total = 356 | 21 (11%) | 0 | 33 (12%) | 0 | 36 (10%) | 4 (3%) |
| RAVLT (T1-immediade recall)  n-total = 254 | 14 (4%) | 0 | 12 (4%) | 0 | 33 (8%) | 0 |
| RAVLT (T5-immediade recall)  n-total = 255 | 0 | 0 | 2 (1%) | 0 | 14 (4%) | 5 (2%) |
| RAVLT (total immediate recall)  n-total = 254 | 9 (4%) | 0 | 9 (4%) | 0 | 28 (7%) | 1 (1%) |
| RAVLT (delayed recall)  n-total = 255 | 1 (1%) | 2 (2%) | 0 | 19 (3%) | 1 (1%) | 6 (5%) |
| RAVLT (delayed correct recognition)  n-total = 251 | 0 | 2 (2%) | 0 | 8 (4%) | 1 (1%) | 11 (5%) |
| RAVLT (delayed false recognition)  n-total = 252 | 4 (3%) | 0 | 16 (5%) | 0 | 19 (5%) | 0 |
| Anxiety (State)  n-total = 466 | 0 | 5 (3%) | 0 | 15 (4%) | 0 | 4 (4%) |
| Anxiety (Trait)  n-total = 466 | 0 | 103 (28%) | 0 | 36 (19%) | 0 | 66 (24%) |
| NEO (N total)  n-total = 462 | 0 | 1 (1%) | 0 | 0 | 5 (1%) | 23 (6%) |
| NEO (E total)  n-total = 461 | 38 (13%) | 0 | 39 (12%) | 0 | 46 (14%) | 0 |
| NEO (O total)  n-total = 462 | 119 (36%) | 0 | 61 (17%) | 0 | 115 (26%) | 0 |
| NEO (A total)  n-total = 463 | 0 | 0 | 0 | 0 | 10 (3%) | 1 (1%) |
| NEO (C total)  n-total = 463 | 8 (4%) | 10 (7%) | 1 (1%) | 8 (5%) | 5 (4%) | 12 (4%) |

Abbreviations: IQ : intelligence quotient; ANT : attention network task; CWI: color-word interference task; RAVLT : Rey auditory verbal learning task; NEO five factor inventory (N: neuroticism, E: extraversion, O: openness, A: agreeableness, C: conscientiousness);
